# Actin filaments act as a size-dependent diffusion barrier around centrosomes

**DOI:** 10.1101/2021.10.21.465377

**Authors:** Hsuan Cheng, Yu-Lin Kao, Lohitaksh Sharma, Wen-Ting Yang, Shih-Han Huang, Hong-Rui Lin, Yao-Shen Huang, Chi-Ling Kao, Lee-Wei Yang, Rachel Bearon, Yu-Chun Lin

## Abstract

The centrosome, a non-membranous organelle, constrains various soluble molecules locally to execute its functions. As the centrosome is surrounded by various dense components, we hypothesized that the centrosome may be bordered by a putative diffusion barrier. After quantitatively measuring the trapping kinetics of soluble proteins of varying size at centrosomes by a chemically inducible diffusion trapping assay, we found that centrosomes were highly accessible to soluble molecules with a Stokes radius of ≤ 5.1 nm, whereas larger molecules rarely reach centrosomes, indicating the existence of a size-dependent diffusion barrier at centrosomes. The permeability of barriers was tightly regulated by branched actin filaments outside of centrosomes. Such barrier gated the microtubule nucleation. We propose that actin filaments spatiotemporally constrain the distribution of molecules at centrosomes in a size-dependent manner.

**Significance Statement:** Centrosome maintains its microenvironment without membrane. Whether the dense protein complexes outside centrosomes including pericentriolar matrix, microtubules, and branched actin filaments provide physical obstruction is unclear yet. We here established a series of new tools for quantitative evaluation of the diffusion rates of varisized soluble proteins in different sub-compartments of centrosomes. Our results demonstrated that branched actin filaments, but not pericentriolar matrix or microtubules, around centrosome have acted as a size-dependent diffusion barrier and physically constrain centrosome microtubule nucleation.

## Introduction

Precise regulation of protein distribution and dynamics ensures proper cellular function and architecture, defects of which are associated with degenerative and neoplastic diseases(1, 2). Whereas active transport requires energy for molecules to move to their destination, soluble molecules diffuse down a concentration gradient without energy expenditure in passive transport(3). To conduct local reactions, cells compartmentalize soluble molecules spatiotemporally by using diffusion barriers in various forms such as nuclear pore complexes, dendritic spine necks, axon initial segments, and ciliary pore complexes(3–6). These passive permeable diffusion barriers serve as conduits between different subcellular compartments and regulate the movement of soluble molecules across adjacent pools. Diffusion barrier deficiency results in protein mislocation/dislocation, which causes several known human diseases(4, 7, 8).

Eukaryotic cells contain endomembrane systems to compartmentalize molecules in membranous organelles for executing different functions(9). Unlike most organelles, the centrosome is a non-membranous organelle; however, it is still capable of assembling hundreds of specific molecules locally and maintaining its microenvironment within the cytosol pool. These attributes ensure its proper architecture and function in terms of microtubule nucleation(10, 11) and formation of the primary cilium, an important protruding structure on the cell surface for sensing extracellular stimuli(10, 12–14). Many scaffold proteins serve as platforms to recruit centrosomal molecules from the cytosolic pool. Moreover, it is well known that the highly dense protein matrix that is present in confined areas is capable of forming permeable diffusion barriers(15). Based on the previous observation that centrosomes are embedded in a cloud of proteins known as the pericentriolar matrix, and emanating microtubules as well as branched actin filaments(16, 17), we hypothesized that a putative diffusion barrier may exist around centrosomes to constrain the distribution and dynamics of centrosomal molecules. To investigate the diffusion behavior of soluble molecules, several conventional approaches including single-particle tracking, fluorescence correlation spectroscopy, fluorescence recovery after photobleaching (FRAP), and tracking photoactivatable fluorescence proteins or microinjected molecules have been widely applied (18–27). However, the diffusion behaviors of molecules uncovered by these conventional methods are determined by both their intrinsic diffusion rates and extrinsic affinity for other immobile scaffolds, which cannot be easily uncoupled. To faithfully uncover the diffusion rates of soluble molecules independently of extrinsic interference, we applied a chemically inducible diffusion trap (CIDT), a previously developed approach, which permits the monitoring of molecule diffusion rates in specific subcellular sites while avoiding interactions between probes and immobile scaffolds in close proximity(3, 18, 28). With the CIDT approach, we have identified a size-dependent diffusion barrier consisting of actin filaments as the key component around centrosomes.

## Results

### Establishment of centrosome-specific CIDT systems

We first describe how to use the CIDT approach to probe diffusion barriers. CIDT in living cells involves three components: rapamycin, a naturally occurring chemical dimerizer; FK506-rapamycin-binding domain (FRB); and immunophilin FK506-binding protein-12 (FKBP)(29). Introduction of rapamycin swiftly dimerizes FRB with nearby molecules of FKBP (**Fig. 1a**)(30). Typically, FRB molecules are tagged with a known targeting sequence (TS in **Fig. 1a**) and consequently anchored to regions of interest (ROIs), whereas FKBP molecules are tagged with diffusive probes (FKBP-probes) that uniformly distribute in the cytosolic pool. When the diffusion probes can access FRB-ROIs, the addition of rapamycin rapidly induces FRB/FKBP dimerization and consequently traps FKBP-probes from the cytosol onto FRB-ROIs (**Fig. 1a**). In the presence of a diffusion barrier, however, the barrier can block certain FKBP-probes from the ROI (**Fig. 1a**). In addition to accessibility, the trapping kinetics of FKBP-probes depend on their diffusion rates across cytosolic pools to FRB-ROIs. By measuring the kinetics of variably sized FKBP-probes that are trapped by FRB-ROIs, putative size-dependent diffusion barriers around FRB-ROIs can be characterized(3, 18). CIDT has been successfully used to identify a size-dependent diffusion barrier at the base of the primary cilium(18), which was also confirmed by other assays(31).

**Figure 1.**
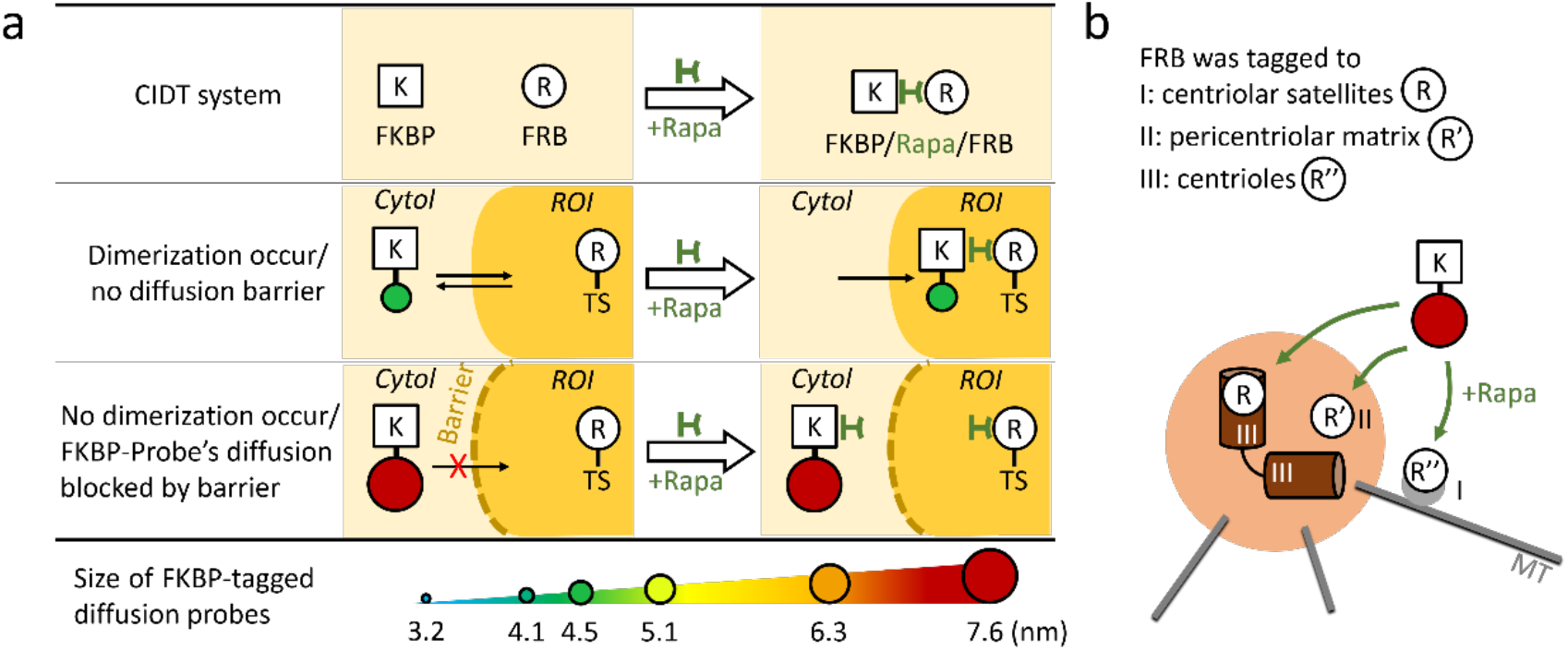
Using the CIDT system to probe a putative centrosomal diffusion barrier. (a) With the CIDT (chemically inducible diffusion trapping) system, YFP-FKBP−tagged diffusive protein probes (FKBP-probes) ranging from 40 to 659 kDa (3.2 nm to 7.6 nm) were used to probe the putative diffusion barrier in cells. FRB was localized to region of interest (ROI) by tagging with the targeting sequence of ROI (TS). Theoretically, in the absence of a barrier (middle panel), probes can freely access and be trapped at ROI via rapamycin (Rapa)-triggered FRB/FKBP dimerization; in the presence of a barrier (lower panel), probe diffusion can be hindered and, thus, dimerization obstructed. (b) To explore the precise location of putative diffusion barriers at centrosomes, FRB (R) was tagged to localize to three different sub-compartments of centrosomes, the centriolar satellites (I), pericentriolar matrix (II), and centrioles (III). The accessibility of FKBP (K)-tagged diffusion probes of various sizes to centrosomes was tested by the CIDT system.

To explore putative diffusion barriers at centrosomes, we constrained FRB molecules at centrosomes by tagging FRB with centrosome-targeting sequences (CTSs) (**Fig. 1b**). It is necessary to select CTSs with low shuttling and highly sustained localization at the designated centrosomal sub-compartments in an attempt to focus on probe diffusion and avoid interference from the active transport of probes, as would be facilitated by CTSs that are highly mobile. Therefore, we took advantage of the FRAP assay to explore and evaluate the dynamics of several previously identified CTS candidates including CEP120C (C terminus of CEP120, which localizes at the centriolar outer wall)(32), CEP170C (C terminus of CEP170, which localizes at subdistal appendages)(33), PACT (C terminus of Pericentrin, which localizes at the pericentriolar matrix), Centrin2 (which localizes to the centriolar lumen)(34), and PCM1F2 (F2 domain of PCM1, which localizes at centriolar satellites)(35). Among these, green fluorescent protein (GFP)-tagged CEP120C and CEP170C showed the most rapid recovery after photobleaching (**Supplementary Fig. 1a-c**) with highly mobile fractions (>20%) and low recovery half-time (<60 sec) (**Supplementary Fig. 1c**), indicating constant centrosome cytoplasmic shuttling of CEP120C and CEP170C. Meanwhile, PCM1F2, PACT, and Centrin2 were capable of constraining a cyan fluorescent protein, Cerulean3 (Ce3), and the FRB molecule in specific sub-compartments of centrosomes (Ce3-FRB-Centrin2 at centrioles; PACT-Ce3-FRB at the pericentriolar matrix; PCM1F2-Ce3-FRB at centriolar satellites; **Supplementary Fig. 2**), show a low shuttling rate with a low mobile fraction (<20% in 5 min) (**Supplementary Fig. 1b-c**), making them suitable CTSs for the CIDT system.

We confirmed that rapamycin treatment triggers the trapping of cytosolic FKBP-tagged probes in the ROIs where the corresponding CTS-FRB proteins reside by co-expressing yellow fluorescent protein (YFP)-tagged FKBP (YFP-FKBP; Stokes radius [Rs], 3.2 nm; molecular weight, 40 kDa) with PCM1F2-Ce3-FRB, PACT-Ce3-FRB, and Ce3-FRB-Centrin2 in U2Os cells, a cell type derived from a human osteosarcoma that exhibits clear centrosomal structures. (**Fig. 2a; Supplementary videos 1−3, left**). The accumulation of YFP-FKBP onto each centrosome locale quickly reached a plateau level, probably due to the full occupancy of FRB binding sites locally (**Fig. 2b**). The half-time (t_1/2_) values for translocation of YFP-FKBP onto centrioles, the pericentriolar matrix, and centriolar satellites were all similar (centrioles: 12.15 ± 2.27 sec; pericentriolar matrix: 13.83 ± 3.09 sec; centriolar satellites: 7.24 ± 0.46 sec; **Table 1**). This confirmed the applicability of the centrosome CIDT method in our subsequent experiments.

**Table 1.** Translocation half-time and probability of varisized probes at centriolar satellites, the pericentriolar matrix, and the centriolar lumen.

| Tested proteins | Stoke radius (nm) | Molecular weight (kDa) | $t_{1/2}$ to CS (sec) | Diffusion probability to CS (%) | $t_{1/2}$ to PM (sec) | Diffusion probability to PM (%) | $t_{1/2}$ to centrioles (sec) | Diffusion probability to centrioles (%) |
| --- | --- | --- | --- | --- | --- | --- | --- | --- |
| YFP-FKBP | 3.2 | 40 | 7.24 ± 0.46 | 99 ± 1% | 13.66 ± 2.38 | 99 ± 1% | 12.15 ± 2.27 | 81 ± 3% |
| YFP-FKBP-Grp1 | 4.1 | 57 | 2.52 ± 1.15 | 99 ± 1% | 19.41 ± 4.19 | 96 ± 2% | 11.74 ± 2.19 | 97 ± 2% |
| YFP-FKBP-Luciferase | 4.5 | 100 | 5.66 ± 2.31 | 99 ± 1% | 21.04 ± 3.10 | 98 ± 2% | 22.19 ± 3.28 | 66 ± 9% |
| YFP-FKBP-Tiam1 | 5.1 | 124 | 9.75 ± 2.39 | 98 ± 2% | 16.84 ± 2.62 | 96 ± 2% | 29.53 ± 4.09 | 96 ± 1% |
| YFP-FKBP-ΔNβ-Gal | 6.3 | 322 | 6.53 ± 3.18 | 76 ± 5% | - | 52 ± 8% | - | 33 ± 6% |
| YFP-FKBP-β-Gal | 7.6 | 659 | 12.41 ± 7.38 | 85 ± 11% | - | 18 ± 3% | - | 10 ± 3% |

\*CS: centriolar satellites; PM: pericentriolar matrix

**Figure 2.**
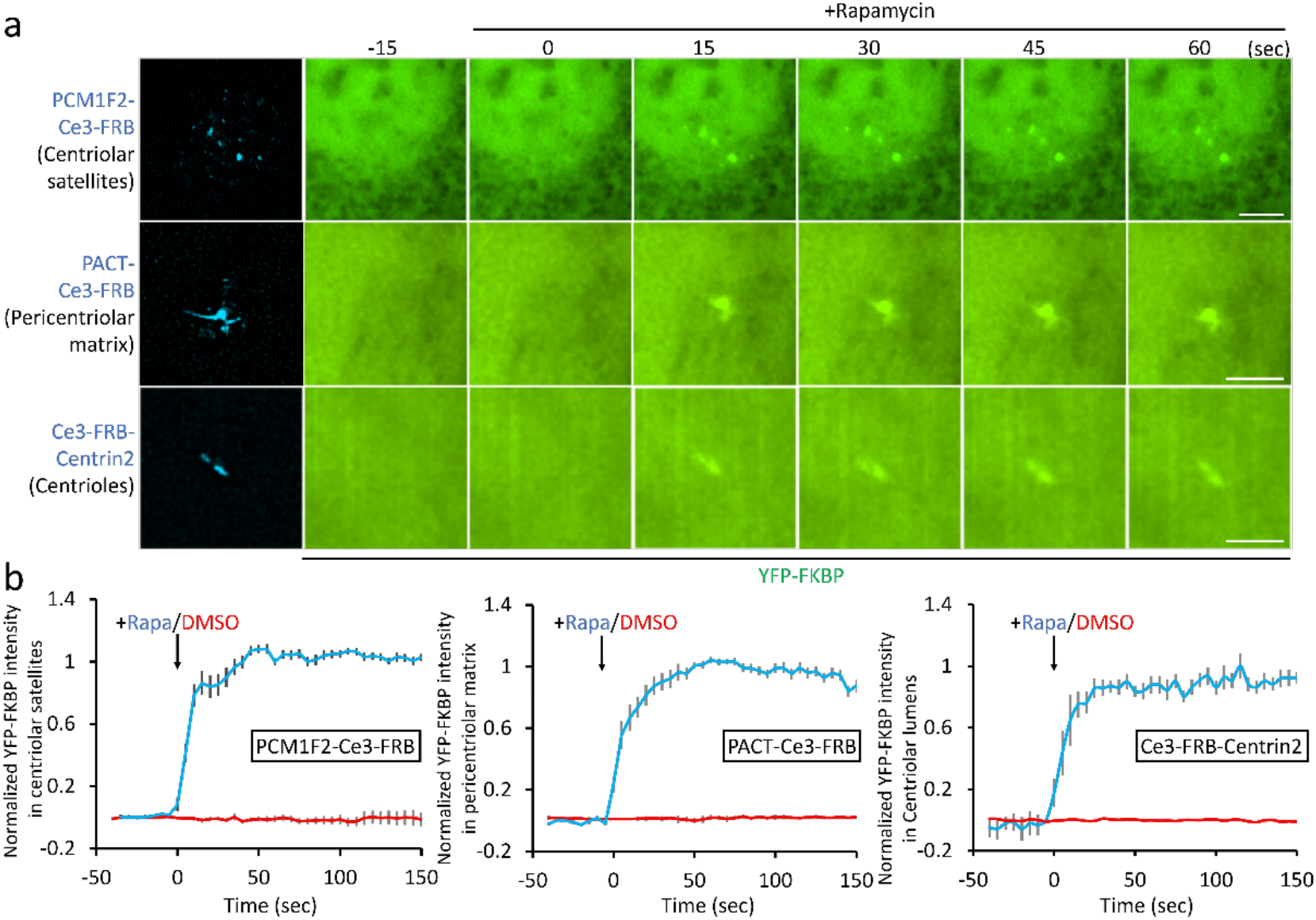
Trapping diffusion probes at three centrosomal sub-compartments. (a) U2Os cells co-transfected with YFP-FKBP and PCM1F2-Ce3-FRB, PACT-Ce3-FRB, or Ce3-FRB-Centrin2 were treated with rapamycin (100 nM). The translocation of YFP-FKBP onto PCM1F2-Ce3-FRB−labeled centriolar satellites, the PACT-Ce3-FRB−labeled pericentriolar matrix, and the Ce3-FRB-Centrin2−labeled centriolar lumen was monitored. Scale bars, 5 µm. (b) The normalized fluorescence intensity of YFP-FKBP accumulation at centrosomes upon rapamycin (100 nM; blue) and DMSO (0.1%, vehicle control; red) treatment. Data are shown as the mean ± S.E.M. The graphs show immediate translocation of YFP-FKBP after rapamycin induction at centriolar satellites (left, n = 24 cells), the pericentriolar matrix (middle, n = 20 cells), and the centriolar lumen (right, n = 23 cells) from seven independent experiments.

### Recruiting varisized diffusion probes onto centrosomes

We investigated the diffusion of varisized proteins at three centrosome locales in U2Os cells. Native Rs (Stoke radius) for YFP-FKBP, YFP-FKBP-Grp1, YFP-FKBP-Luciferase, YFP-FKBP-Tiam1, YFP-FKBP-ΔNβ-Gal, and YFP-FKBP-β-Gal were 3.2, 4.1, 4.5, 5.1, 6.3, and 7.6 nm, with molecular weights of 40, 57, 100, 124, 322, and 659 kDa, respectively, according to previous measurements (**Table 1**)(18). We next co-expressed PCM1F2-Ce3-FRB and each of the YFP-FKBP−tagged probes in U2Os cells. All varisized diffusion probes immediately translocated onto the centrosome peripheral, centriolar satellites, within 13 sec upon rapamycin treatment (**Fig. 3a,b, left, Supplementary Fig. 3, Table 1, and Supplementary video 1**). Interestingly, whereas probes with an Rs as large as 5.1 nm still showed rapid translocation onto all three centrosomal sub-compartments, YFP-FKBP-ΔNβ-Gal (6.3 nm) and YFP-FKBP-β-Gal (7.6 nm) had not been recruited onto the pericentriolar matrix and centriolar lumen by 5 min (**Fig. 3a,b, middle and right; Supplementary Figs. 4,5, Table 1, and Supplementary videos 2,3**).

**Figure 3.**
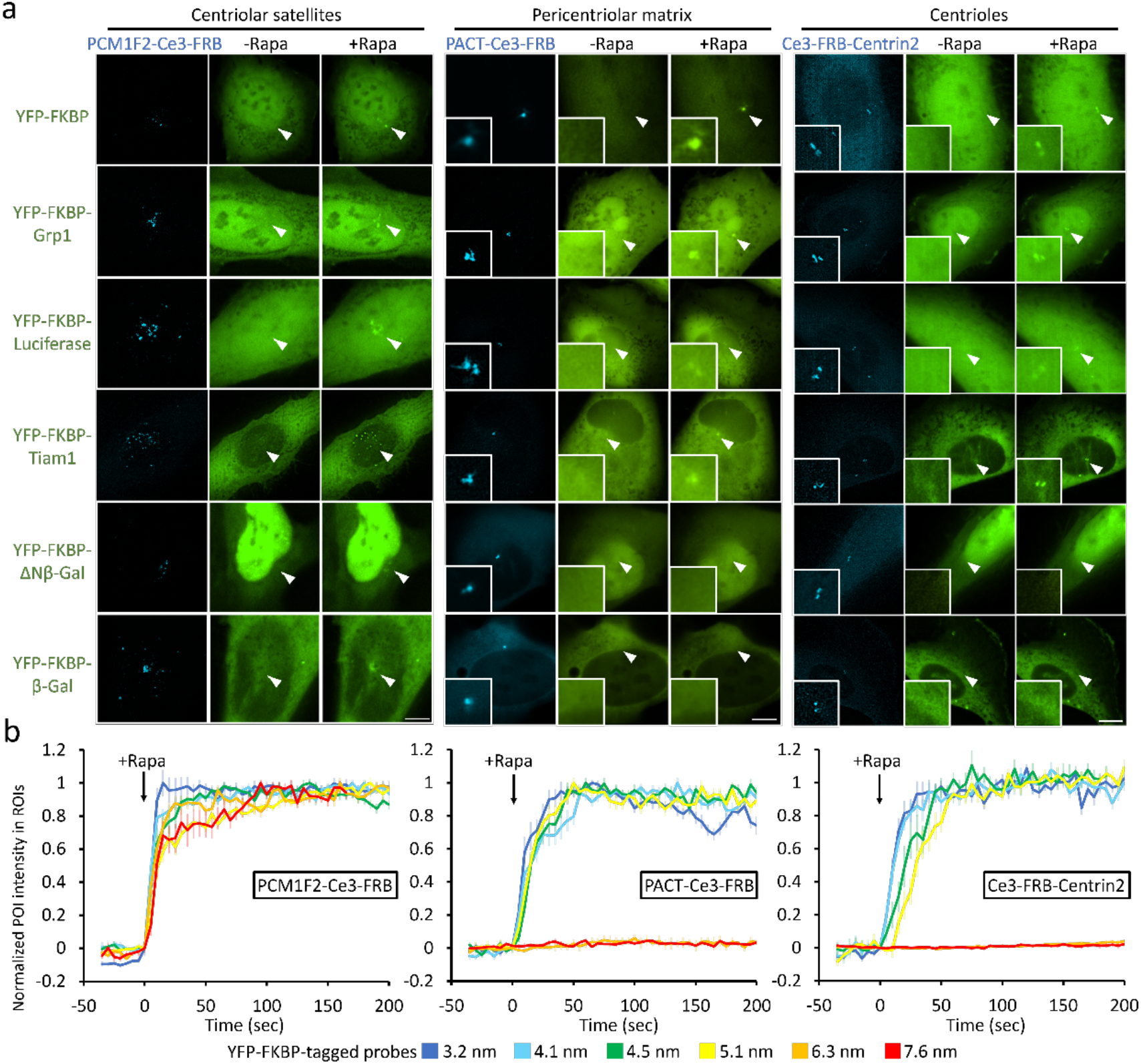
Size-dependent accessibility of diffusion probes into centrosomes. (a) U2Os cells co-transfected with each YFP-FKBP−labeled varisized probe (YFP-FKBP, YFP-FKBP-Grp1, YFP-FKBP-Luciferase, YFP-FKBP-Tiam1, YFP-FKBP-ΔNβ-Gal, and YFP-FKBP-β-Gal) and Ce3-FRB with labels specific for centrosomal sub-compartments were treated with rapamycin (100 nM). Arrowheads indicate sites of centrosomes. Insets show higher-magnification images of the centrosomal regions. Scale bars, 10 µm. (b) The normalized fluorescence intensity of each probe at centriolar satellites (left, n = 89 cells), pericentriolar matrix (middle, n = 98 cells), and centriolar lumen (right, n = 75 cells) from three independent experiments. Data are shown as the mean ± S.E.M.

We further quantified the translocation probability by extending the translocation time to 2 hr after trapping the diffusive probes by rapamycin treatment. Cells transfected with 18 combinations of one CFP-FRB-ROI and one YFP-FKBP−tagged probe representing each condition were treated with rapamycin for 2 hr (**Table 1**). The rapamycin groups demonstrated a tendency to decline in their translocation rate as the size of the diffusive probes increased. The translocation rates were high and the differences were subtle between probes with Rs of ≤ 5.1 nm at the periphery of the centrosome (centriolar satellites). However, the change in translocation rates was especially drastic at the core centrosomal regions (pericentriolar matrix and centrioles) for probes with Rs ≥ 6.3 nm. The translocation probability of YFP-FKBP-ΔNβ-Gal (Rs, 6.3 nm) at the pericentriolar matrix and centrioles showed a nearly twofold change, similar for YFP-β-Gal (Rs, 7.6 nm), which had an even lower translocation rate of 10 ± 3% at the centrioles after a 2-hr rapamycin treatment (**Table 1**).

In conclusion, the diffusion kinetics of soluble proteins ranging from an Rs of 3.2 to 7.6 nm (40−659 kDa) at centriolar satellites were similar, whereas their diffusion/accumulation at the pericentriolar matrix and centrioles showed a negative correlation. The probability of accessing peripheral centriolar satellites for soluble diffusive probes with Rs ≥ 6.3 nm (322 kDa) is still high but plummets at the pericentriolar matrix and centrioles. The poor accessibility of large probes to the centrosome core is not determined by their diffusivity in the cytosol, as the trapping kinetics of those probes to centriolar satellites and to the plasma membrane are all rapid (**Fig. 3, Supplementary Figs. 3,6; Table 1; Supplementary video 1**). Given that the rapid recruitment occurred within a minute, the mTOR pathway was not likely to contribute to the diffusive effect despite the addition of rapamycin. Taken together, these results revealed the existence of a size-dependent diffusion barrier between the core centrosomes and cytosol pool.

### Actin filaments contribute to the centrosomal diffusion barrier

We next aimed to explore the molecular components of the centrosomal diffusion barriers. Centrosomes are known to be microtubule- and actin-organizing centers that are surrounded by these cytoskeletal elements(10, 11, 36–38). We hypothesized that either microtubules or actin filaments contribute to the function and composition of centrosomal diffusion barriers, as it is known that the cytoskeleton not only provides mechanical support but also serves as a critical component of permeable passive barriers in cells(39).

To examine the role of microtubules in centrosomal diffusion barriers, U2Os cells were treated with Nocodazole to depolymerize the cellular microtubule polymers to soluble tubulins(40). An immunostaining assay confirmed that microtubule depolymerization occurred 10−20 min after Nocodazole treatment (**Fig. 4a**). We then carried out the CIDT assay on cells with either intact or depolymerized microtubules. Live-cell imaging showed that YFP-FKBP-ΔNβ-Gal was blocked from entering centrioles even after microtubule depolymerization, indicating that the contribution of microtubules with respect to the diffusion barrier was rather trivial (**Fig. 4b,c**).

**Figure 4.**
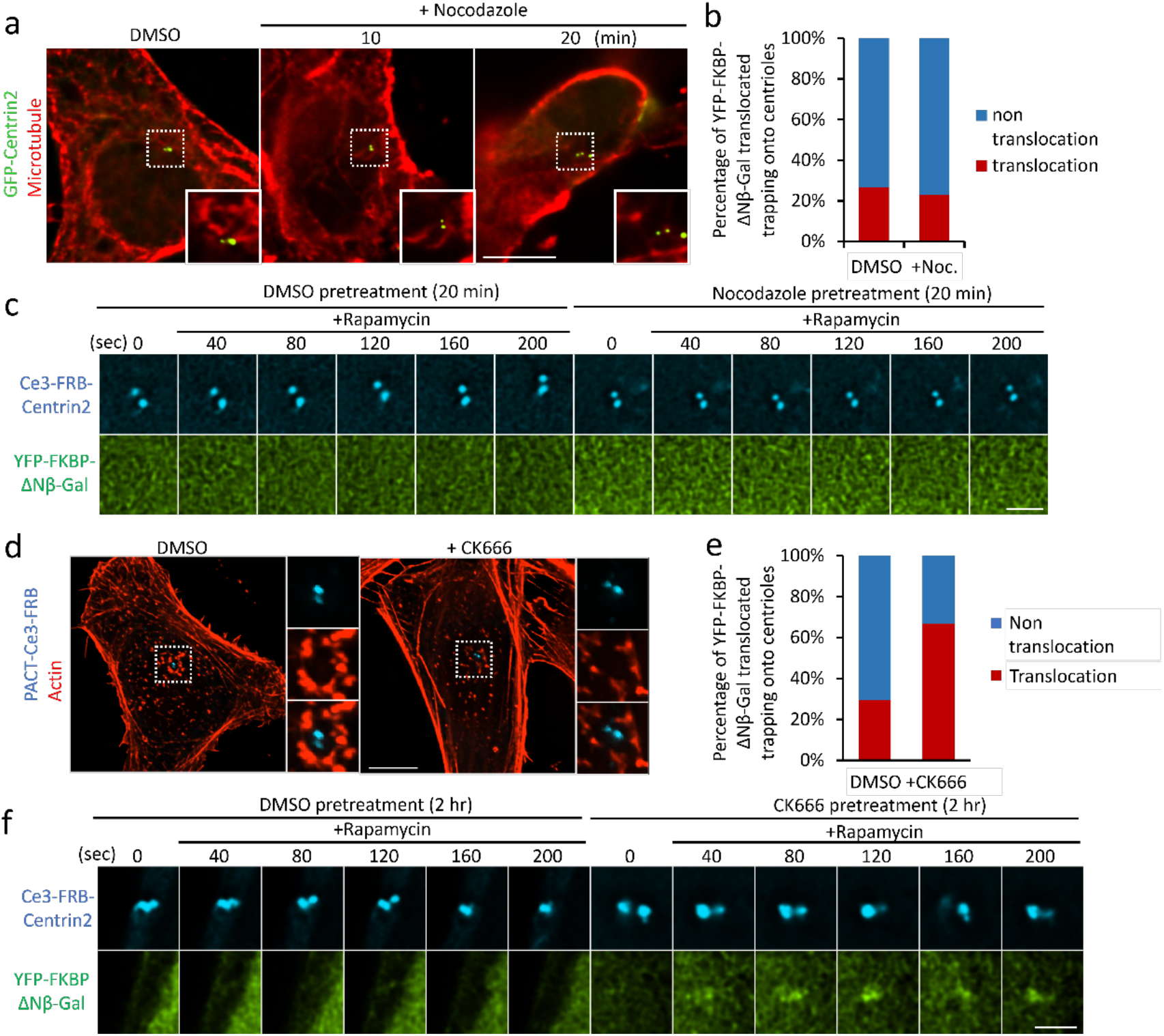
Branched actin filaments are key components of centrosomal diffusion barriers. (a) Depolymerization of microtubules by nocodazole (10 µM) treatment. U2Os cells that expressed GFP-Centrin2 (green, a marker of centrosomes) were incubated with antibodies against α-tubulin to label microtubules (red) in the absence (0.1% DMSO) or presence of nocodazole treatment for 10 and 20 min. Insets show higher-magnification images of the centrosomal regions. Scale bar, 10 µm. (b) The percentage of DMSO- or nocodazole-treated cells exhibiting YFP-FKBP-ΔNβ-Gal translocation (red; n = 26 cells). (c) U2Os cells co-transfected with YFP-FKBP-ΔNβ-Gal and Ce3-FRB-Centrin2 were treated with rapamycin (100 nM) prior to and after microtubule depolymerization with nocodazole. Scale bar, 2 µm. (d) U2Os cells expressing PACT-Ce3-FRB (blue, a marker of centrosomes) were incubated with phalloidin to label the actin filaments (red) with DMSO (0.1%) or CK666 (0.4 mM) for 2 hr. Right panels show the higher-magnification images of the centrosomal regions. Scale bar, 10 µm. (e) The percentage of DMSO- or CK666-treated cells exhibiting YFP-FKBP-ΔNβ-Gal translocation onto centrioles (red; n = 15 cells). (f) U2Os cells co-transfected with YFP-FKBP-ΔNβ-Gal and Ce3-FRB-Centrin2 were treated with 100 nM rapamycin in the absence and presence of actin depolymerization by 0.4 mM CK666 incubation for 2 hr. Scale bar, 2 µm.

We next investigated the role of actin filaments. Centrosomes are surrounded by Arp2/3-associated branched actin-filaments(41). We disrupted actin filaments in U2Os cells by the addition of CK666, an Arp2/3 inhibitor(42, 43), which perturbs actin polymerization(44). Immunofluorescence results showed the reduction in actin filaments around the centrosomes after 2 hr of CK666 treatment (**Fig. 4d**). A CIDT assay in the presence of CK666 showed that the decrease in centrosomal actin filaments allowed the entry of YFP-FKBP-ΔNβ-Gal, which could not access the centriolar lumen in the absence of CK666 (**Fig. 4e,f**), indicating the substantial contribution of actin filaments to the composition of the centrosomal size-dependent diffusion barrier.

### The permeability of centrosomal diffusion barriers decreases in anaphase

At anaphase, an increase in the amount of branched actin around centrosomes was observed(45). We therefore tried to characterize the permeability of centrosomal diffusion barriers in anaphase cells. In interphase HeLa cells, YFP-FKBP-Luciferase could access PACT-labeled pericentriolar matrix (**Fig. 5; Supplementary Video 4**). However, the entry of YFP-FKBP-Luciferase onto centrosomes was blocked in anaphase cells, indicating the permeability of centrosomal diffusion barriers temporally decreased during anaphase (**Fig. 5; Supplementary Video 4**).

**Figure 5.**
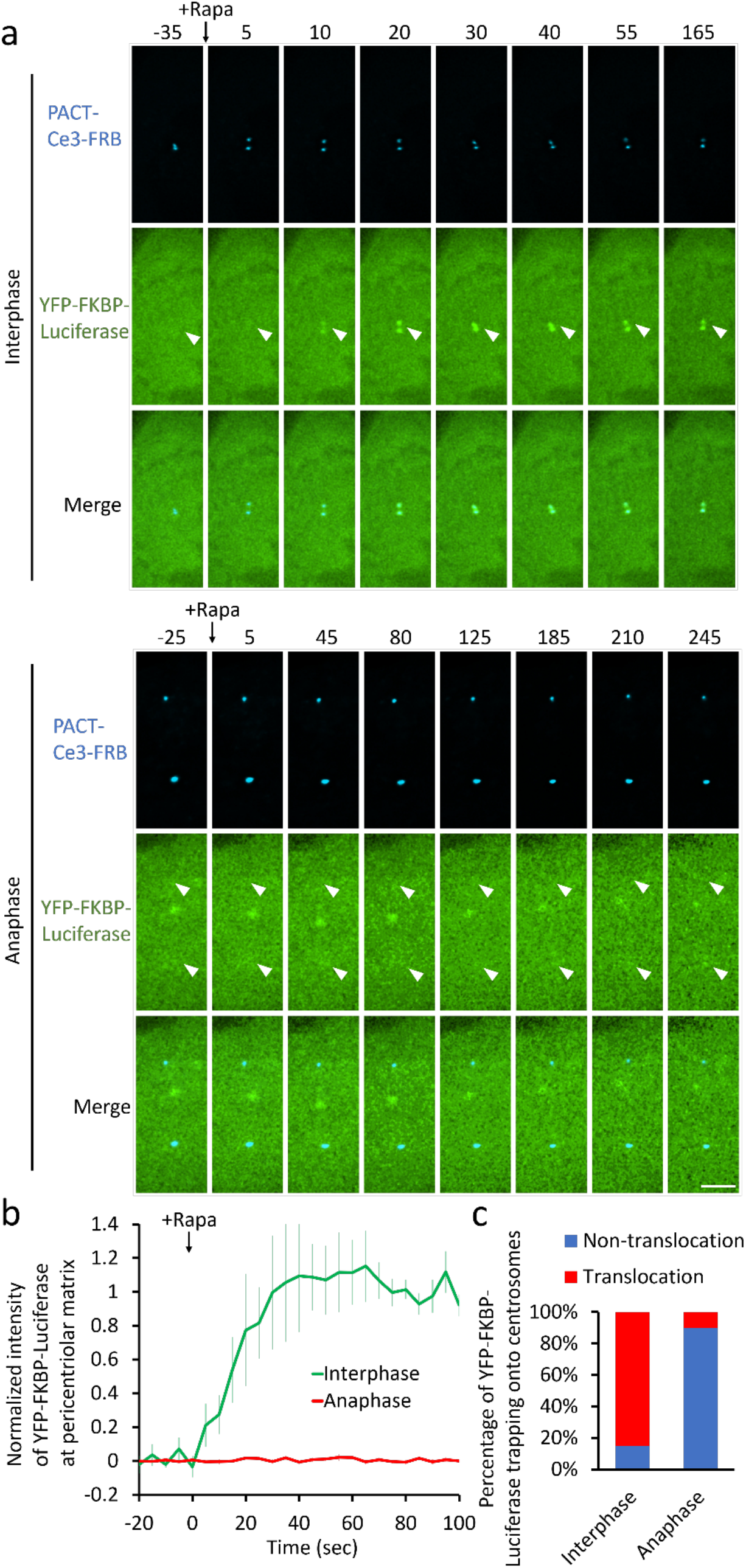
The permeability of centrosomal diffusion barriers decreases in anaphase. (a) HeLa cells co-transfected with PACT-Ce3-FRB and YFP-FKBP-Luciferase after (anaphase cells) or without (interphase cells) synchronization. Transfected cells in interphase (upper panel) or anaphase (lower panel) were treated with 100 nM rapamycin (Rapa). Arrowheads indicate sites of centrosomes in the YFP channel. Scale bar, 5 µm. (b) The normalized fluorescence intensity of YFP-FKBP-Luciferase accumulation at centrosomes in interphase (green, n= 12 cells) or anaphase (red, n=10 cells) cells upon rapamycin (100 nM) treatment. Data are shown as the mean ± S.E.M. (c) The percentage of interphase or anaphase cells exhibiting YFP-FKBP-Luciferase translocation.

### The actin-based diffusion barrier physically regulates *r*-TuRC and microtubule nucleation

The centrosome acts as a microtubule organizing center via recruiting the *r*-tubulin ring complex (*r*-TuRC), a key protein complex (Rs: ∼15 nm)(46) required for microtubule nucleation, onto centrosome(47). We tried to examine whether the centrosome diffusion barrier gates the recruitment of *r*-TuRC at centrosomes. The density of one major component in *r*-TuRC, *r*-tubulin, in control and CK666-treated centrosomes was labeled and quantified. Disruption of actin-based diffusion barriers significantly increased the amount of *r*-tubulin at centrosomes (**Fig. 6a,b**). We next transfected U2Os cells with EB1-mNeon, a microtubule plus-end marker tagged with a green/yellow fluorescent protein, mNeon, to track microtubule growth (25 µm in diameter) in the cells with either an intact or disrupted (CK666 treated) diffusion barrier (**Fig. 6c**). We quantitatively measured the frequency of microtubule nucleation and the microtubule elongation rate, events indicative of microtubule growth in the centrosome core region and in the cytosol, respectively. The number of nascent microtubules derived from centrosomes was significantly increased in the absence of a diffusion barrier as compared with that in control cells (**Fig. 6d**). Disruption of the diffusion barrier did not, however, affect the elongation rate of microtubules in cytosol (**Fig. 6e**). These results suggest that recruitment of *r*-TuRC and initiation of microtubule growth around centrosomes were hindered by the diffusion barrier, whereas their elongation rate was not perturbed after they had elongated away from the centrosome.

**Figure 6.**
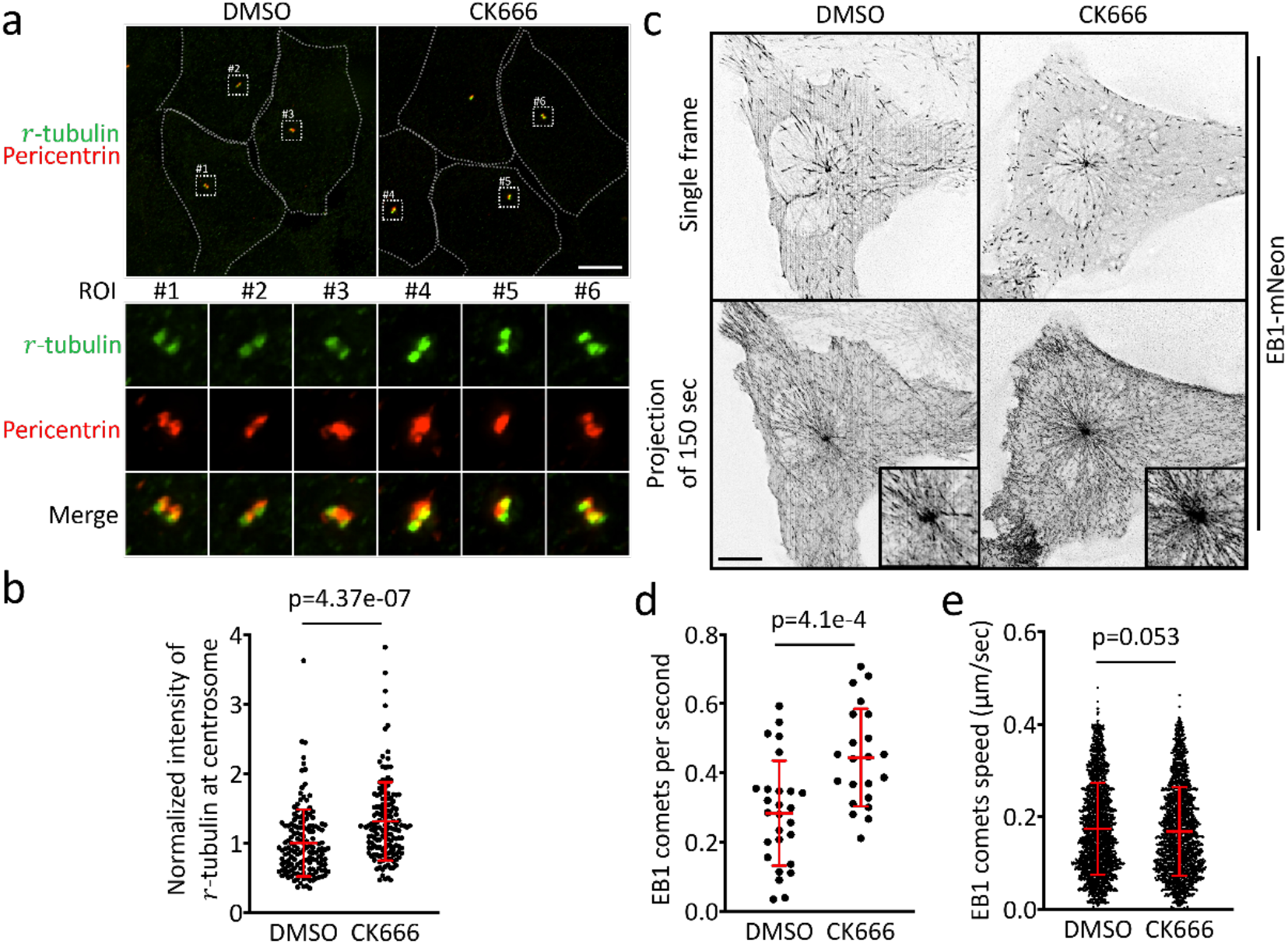
Centrosomal diffusion barriers constrain *r*-TuRC recruitment and microtubule nucleation. (a) U2Os cells were incubated with antibodies against *r*-tubulin (green; a marker of *r*-TuRC) or pericentrin (red; a marker of centrosomes) in the absence (0.1% DMSO) or presence of CK666 treatment (0.4 mM) for 2 hr. The lower panel shows higher-magnification images of the indicated region of interest (ROI). Scale bar, 10 µm. (b) The normalized intensity of *r*-tubulin at centrosome in control (0.1% DMSO, 2 hr) or CK666 (0.4 mM, 2 hr)-treated cells. Individual data points and the mean ± SD (in red) are shown (n=150 and 150 cells in DMSO and CK666-treated group, respectively; four independent experiments). (c) U2Os cells expressing EB1-mNeon were incubated with 0.1% DMSO or 0.4 mM CK666 for 2 hr. Images were captured over 150 sec and were overlaid to show the overall tracks of the microtubules. The insets show enlarged images of the centrosomal regions. Scale bar, 10 μm. (b,c) The number of microtubule tracks that emanate from centrosomes (d) and the microtubule elongation rate (e) in the absence and presence of CK666 treatment (0.4 mM). Individual data points and the mean ± SD (in red) are shown (n = 27 and 22 cells in the DMSO- and CK666-treated groups, respectively; three independent experiments).

## Discussion

We here established a centrosome-specific CIDT system that enables rapid recruitment of varisized cytosolic diffusive probes to three centrosome locales, ranging from the core to the periphery. Diffusive probes with an Rs of ≤ 5.1 nm (124 kDa) can freely access all centrosome sub-compartments, whereas the accessibility of large diffusive probes was hindered. Probes with an Rs of ≥ 6.3 nm (322 kDa) could not reach the pericentriolar matrix and centrioles, indicating the existence of a size-dependent diffusion barrier outside the centrosomal core region. The centrosomal diffusion barrier was perturbed after disruption of branched actin filaments. Disruption of the centrosomal diffusion barrier enabled the entry of large molecules and *r*-TuRC as well as increased microtubule nucleation. The permeability of actin-based barriers was temporally decrease in anaphase. In summary, these results suggest that actin-based diffusion barriers outside of centrosomes spatiotemporally gate the diffusion of molecules in a size-dependent manner.

According to our CIDT results, cytosolic molecules with an Rs of ≤ 5.1 nm can freely access centrosomal core regions. Therefore, most cytosolic components, such as small proteins, metabolites, ions, mRNAs, and others, are able to reach the interior of centrosomes. Intuitively, anchoring via their affinity for centrosomal scaffold proteins should be the major mechanism by which small molecules are retained at centrosomes for local reactions. Conversely, without active transport mediated by motor proteins, large molecules (Rs ≥ 6.3 nm; 322 kDa) such as protein complexes, aggresomes, and ribosomes are prohibited from centrosomes. For example, proteasomes are soluble protein complexes that locally trigger proteolytic degradation at centrosomes (48). Because of their very large size (20 S proteasome: ∼750 kDa)(49), we assume that proteasomes cannot access centrosomal core region to directly trigger protein degradation. Alternatively, centrosomal proteins may diffuse to the cytosol pool that is proximal to centrosomes for proteasome-mediated degradation. Indeed, excess centrosomal proteins and misfolded proteins accumulate in regions proximal to centrosomes upon inhibition of proteasome activity, which supports our assumption(48, 50).

We demonstrated that branched actin filaments act as a diffusion barrier around centrosomes. Branched actin filaments are also abundant in dendritic spines, the axonal initial segment, and other subcellular regions (51–53). Not surprisingly, the dense actin meshwork in these sites also functions as a diffusion barrier. Actin filaments in dendritic spines restrict the mobility of proteins larger than 100 kDa. This restriction is reduced during actin reformation in structural plasticity or after actin filaments are severed by latrunculin A (46). The actin-based diffusion barrier in axonal initial segments ensures axon integrity and neuron cell polarity by preventing dendric protein−positive vesicles from accessing the axonal lumen (54). As centrosomes are transformed into basal bodies at the entrance to the lumen of the primary cilium during G0 phase, we assumed that centrosomal diffusion barriers may also halt the entry of cytosolic molecules into the ciliary lumen. However, previous studies demonstrated that proteins with an Rs as large as 7.9 nm can access the ciliary lumen (18, 31). One possible explanation for this is that centrosomal diffusion barriers do not span the entire ciliary base. Ultrastructural observations of branched actin filaments and the ciliary base are needed to confirm this assumption.

Recently, the discovery of phase-separation mechanisms in cells has provided new insights into how non-membrane-bound components assemble and maintain their structures(55, 56). The organelles with phase transition properties generate a gel-like microenvironment in which to concentrate centrosomal molecules locally and to separate this compartment physically from the cytosol(57). However, whether this centrosomal condensate results in any restrictions on solute mobility is still unclear. According to our CIDT results, the accessibility of molecules at centrioles and the pericentriolar matrix is similar, indicating that cytosol molecules smaller than 124 kDa can freely diffuse across the pericentriolar matrix and reach the centriolar wall. Moreover, disruption of branched actin filaments allows the trapping of YFP-FKBP-ΔNβ-Gal (320 kDa) at centrioles. These results indicated that the pericentriolar matrix does not hinder the diffusion of molecules as large as 124 kDa.

Previous studies demonstrated that centrosomal branched actin acts as a negative regulator of microtubule nucleation(45, 58). However, how branched actin at the centrosome suppresses microtubule nucleation is still unclear. Our results indicated that branched actin prevented large molecules from crossing the centrosome boundary. It is thus plausible that microtubule nucleation would also be physically hindered, although microtubule elongation, which occurs outside of the centrosomes, was not affected by diffusion barriers.

The approaches developed here have powerful applications beyond probing diffusion mechanisms. Chemically inducible dimerization systems have been widely used to spatiotemporally manipulate cellular architecture and signaling by recruiting proteins of interest (Rs of ≤ 5.1 nm) onto target sites in cells (18, 59–61). The current study has established a new system that enables rapid recruitment of proteins of interest onto the centrioles, pericentriolar matrix, or centriolar satellites within seconds. It offers a feasible way to spatiotemporally manipulate molecular signaling at three different centrosomal sub-compartments. This approach should allow us to address previously intractable questions in centrosome biology, a discipline that has a strong translational link to a variety of centrosome-related diseases.

## Methods and materials

### Cell Culture and Transfection

U2Os cells and HeLa cells (ATCC) were maintained at 37°C in 5% CO_2_ in DMEM (Corning) supplemented with 10% fetal bovine serum (Gibco), penicillin, and streptomycin (Corning). Twenty-four hours before live-cell imaging or drug treatment, plasmid DNA transfection was performed with the LT-X transfection reagent (Invitrogen). Transfected cells were plated on poly-D-lysine (Sigma-Aldrich)−coated borosilicate cover glass (Paul Marienfeld) in six-well culture plates (Sigma-Aldrich).

### Cell synchronization

HeLa cells were transfected with YFP-FKBP-Luciferase and PACT-Ce3-FRB plasmids using Fugene HD transfection reagent (Promega) 20-24 hr before synchronization. After transfection, cells were incubated with 2 mM thymidine (Sigma) in supplemented DMEM (Corning) for 16-18 hr to arrest cells at G1/S phase, followed by 12 hr of 2.5 ng/mL RO3306 (Sigma) incubation in medium to block the cells at G2/M phase. To enrich the population of anaphase cells, cells were washed and released to RO3306-free medium for 50-60 mins before imaging.

### Image Acquisition

Live-cell imaging was carried out with a Nikon T1 inverted fluorescence microscope with a 60× or 100× oil objective (Nikon), DS-Qi2 CMOS camera (Nikon), and 37°C, 5% CO_2_ heated stage (Live Cell Instrument). Images with multiple *z*-stacks were processed with Huygens Deconvolution software (Scientific Volume Imaging). Image analysis and the maximum intensity projections of images were generated with Nikon Elements AR software.

### Fluorescence recovery after photobleaching assay

U2Os cells transfected with the indicated constructs (GFP-Centrin2, GFP-CEP120C, GFP-CEP170C1, PCM1F2-GFP, or PACT-GFP) were imaged every 10 sec for 10 min on a confocal laser scanning microscope (Zeiss LSM780). The centrosomal regions of cells expressing the indicated constructs were photobleached and allowed to recover for 300 sec. The time-lapse background-subtracted fluorescence intensity for GFP was quantified with Nikon Elements AR software.

### Measurement of the kinetics of soluble proteins translocating onto centrosomes

U2Os cells were transfected with the indicated constructs as described above. Cells transfected with YFP-FKBP−tagged diffusion probes and Ce3-FRB−tagged CTSs were imaged every 5 sec for 4 min on a Nikon T1 inverted fluorescence microscope with a 60× oil objective (Nikon), DS-Qi2 CMOS camera, and 37°C, 5% CO_2_ heated stage. We added 100 nM rapamycin (LC Laboratories) during imaging. Time-lapse images were processed by Huygens Deconvolution software. The intensity of YFP-FKBP-tagged proteins at centrosomes upon rapamycin treatment was measured by Nikon Elements AR software.

### Immunofluorescence staining

Cells were plated on poly-D-lysine−coated borosilicate glass Lab-Tek eight-well chambers (Thermo Scientific). For microtubule immunofluorescence labeling, cells were fixed for 15 min at room temperature in 4% paraformaldehyde (Electron Microscopy Sciences). Fixed samples were permeabilized with 0.1% Triton X-100 (Sigma-Aldrich) in phosphate-buffered saline (PBS) for 40 min. Cells were gently rinsed before being incubated in blocking solution (2% bovine serum albumin in PBS) for 30 min at room temperature. Primary antibodies against α-tubulin (1:1000 dilution; T6199, Sigma-Aldrich), diluted in blocking solution, were used to stain the cells for 1 hr at room temperature. Secondary antibodies diluted in blocking solution (1:1000 dilution) were incubated with cells for 1 hr at room temperature followed by a gentle rinse with PBS.

For immunofluorescence labeling of actin filaments, cells transfected with PACT-Ce3-FRB were seeded on poly-D-lysine−coated borosilicate glass Lab-Tek eight-well chambers. Cells were rinsed with pre-warmed PBS before being incubated in 4% paraformaldehyde (Electron Microscopy Sciences) dissolved in cytoskeleton-preserving buffer (80 mM PIPES, pH 6.8; 5 mM EGTA; 2 mM MgCl_2_)(62). Cells were permeabilized with 0.5% Triton X-100 (in PBS) followed by Phalloidin 594 (1:20 dilution in PBS; A12381, Invitrogen) staining for 1 hr at room temperature.

### Tracking microtubule growth

U2Os cells were transfected with EB1-mNeon and seeded on poly-D-lysine−coated borosilicate cover glass in six-well culture plates (Sigma-Aldrich) 24 hr prior to imaging. Live-cell imaging was acquired using a 5-sec interval over a duration of 5 min and three *z*-stacks. Images were processed with Huygens Deconvolution software and rolling ball correction (Nikon NIS Elements) before being analyzed with ImageJ (Fiji) TrackMate.

### Statistical analysis

Statistical analysis was performed with an unpaired two-tailed Student’s t-test. An F-test was used to determine whether variances were equal or not. P < 0.05 indicates a significant difference. P < 0.01 indicates a highly significant difference.

## ACKNOWLEDGEMENTS

The original pilot study was conducted by Y.C.L. in Dr. Takanari Inoue’s lab (Johns Hopkins University). This study was supported by the Ministry of Science and Technology (MOST), Taiwan (MOST grant numbers 109-2636-B-007-003 and 108-2638-B-010-001-MY2 to Y.C.L.).

## Author Contributions

H.C., Y.L.K, and Y.C.L. designed the experiments. H.C., Y.L.K., Y.S.H., and Y.C.L. conducted the experiments. H.C. and W.T.Y. conducted FRAP experiments. S.H.H conducted cell synchronization experiments. L.S., H.R.L., L.W.Y., and R.B analyzed the translocation kinetics. H.C., Y.L.K., C.L.K., R.B., and Y.C.L. wrote the manuscript.

The authors declare no conflict of interest.

## Figure

**Supplementary Figure 1.**
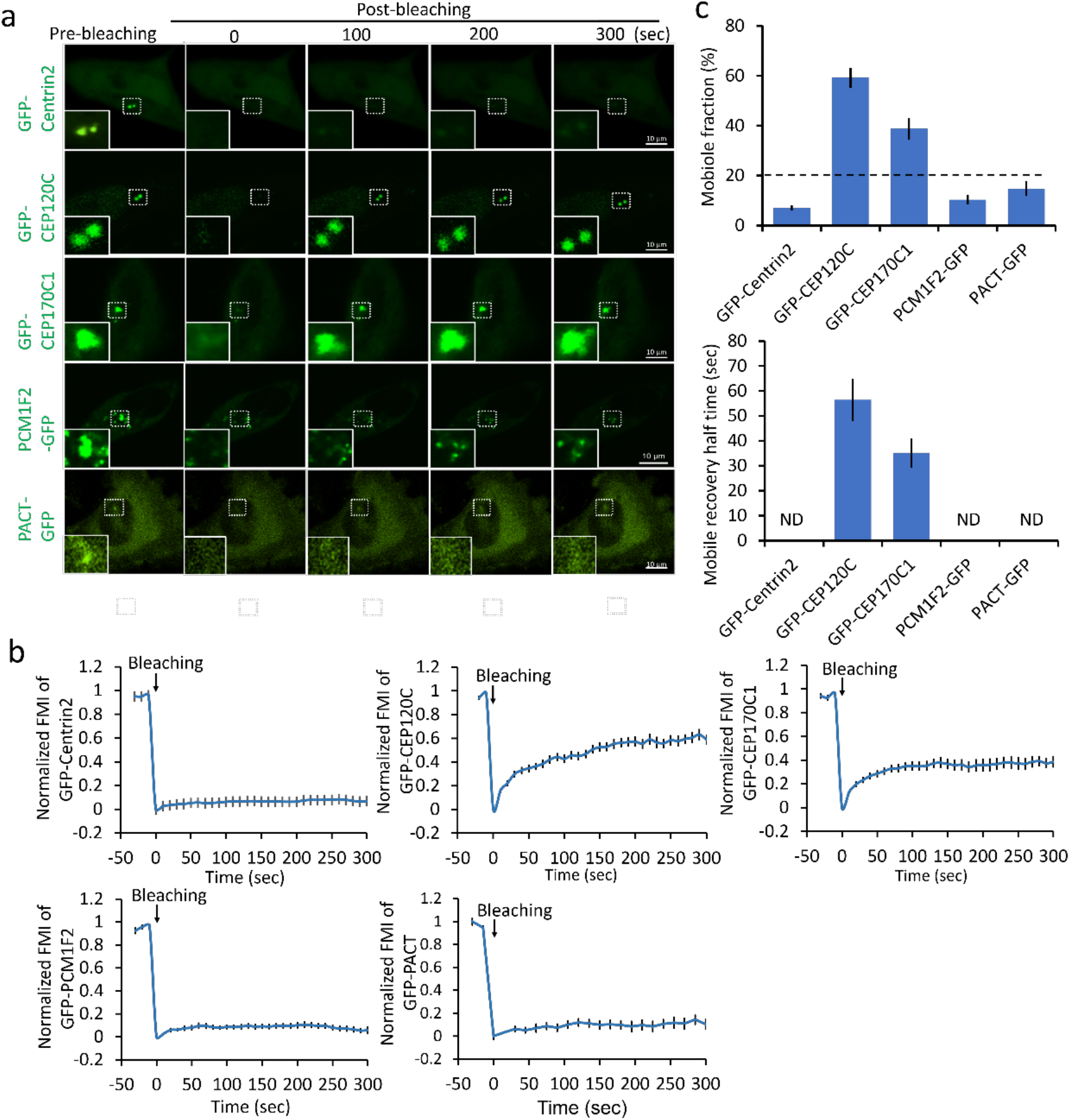
FRAP assay showing low shuttling rate of GFP-Centrin2, PCM1F2-GFP, and PACT-GFP at centrosomes. (a) U2Os cells transfected with centrosome-targeting proteins GFP-Centrin2, GFP-CEP120C, GFP-CEP170C1, PCM1F2-GFP, and PACT-GFP were observed for 300 s after photobleaching. Images of the fluorescently labeled centrosomes are shown. Scale bars: 10 µm. (b) Graphs showing recovery of each GFP-tagged centrosome-targeting protein by normalized fluorescence mean intensity (mean ± S.E.M). (c) The mobile fraction percentage (top) and mobile recovery half-time (bottom) of GFP-Centrin2, GFP-CEP120C, GFP-CEP170C1, PCM1F2-GFP, and PACT-GFP (mean ± SEM; n = 13, 21, 14, 13, and 18, respectively; three to five independent experiments).

**Supplementary Figure 2.**
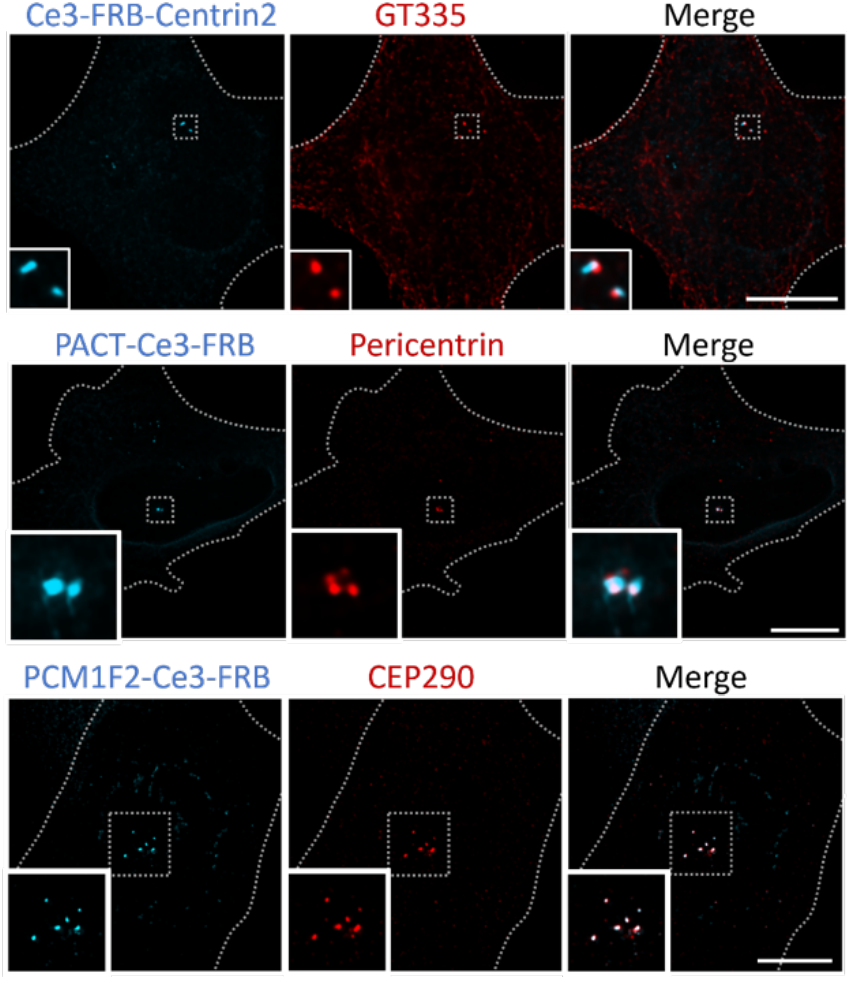
The distribution of Ce3-FRB-Centrin2, PACT-Ce3-FRB, and PCM1F2-Ce3-FRB. Cells transfected with Ce3-FRB-Centrin2, PACT-Ce3-FRB, or PCM1F2-Ce3-FRB were fixed and labeled with anti-GT335 (a marker of centrioles), anti-Pericentrin (a marker of pericentriolar matrix), and anti-CEP290 (a marker of centriolar satellites), respectively. The insets show enlarged images of each centrosomal region. Scale bars: 10 µm.

**Supplementary Figure 3.**
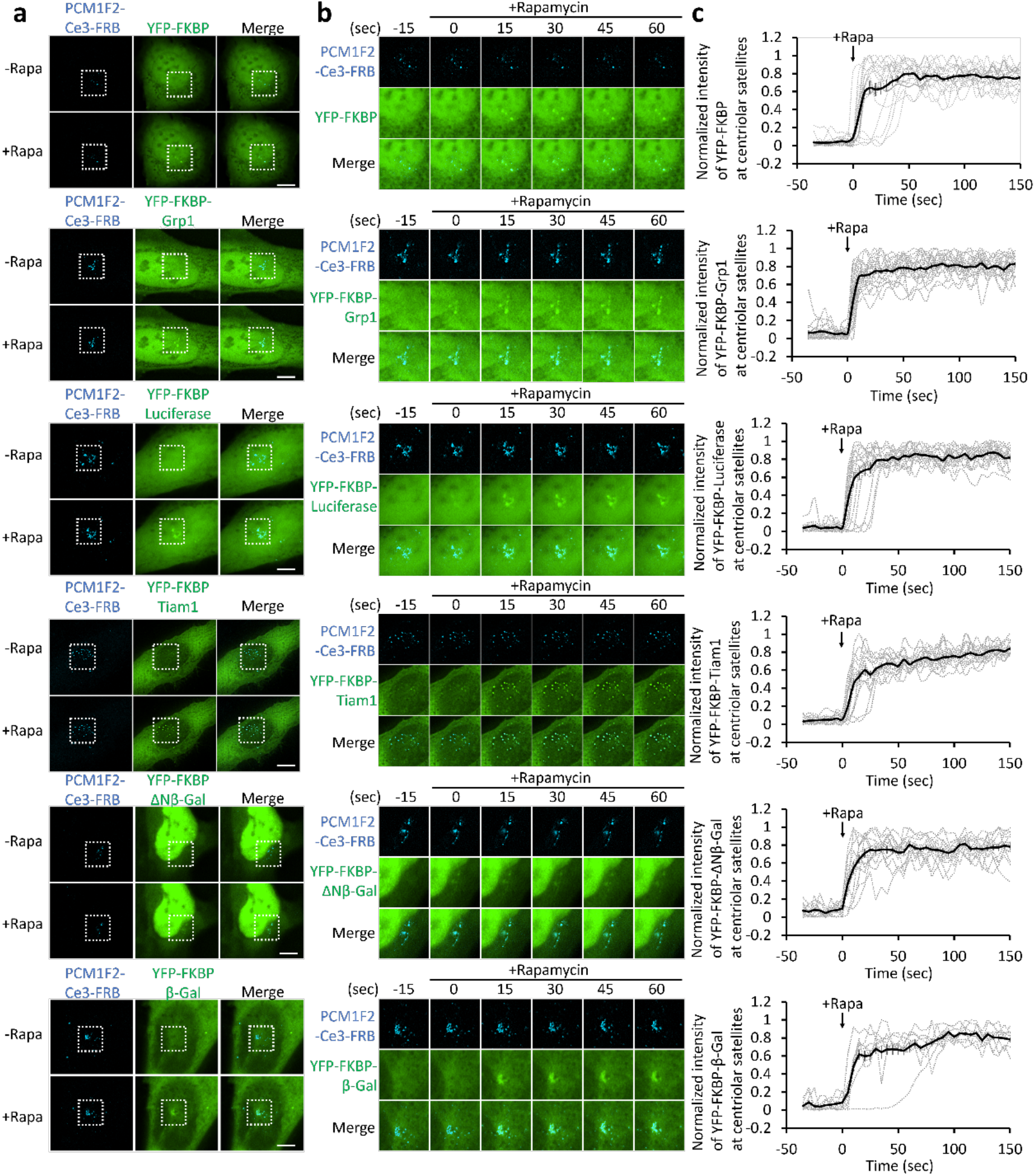
Accessibility of diffusion probes to centriolar satellites. (a) U2Os cells co-transfected with PCM1F2-Ce3-FRB and each YFP-FKBP−tagged varisized probe were treated with rapamycin (100 nM). The dotted line box indicates the centriolar satellite region. Scale bars: 10 µm. (b) Video frames from a movie of the centriolar satellite region as indicated by the dotted line boxes in A upon rapamycin treatment. (c) The normalized intensity of YFP-FKBP−tagged diffusive probes at PCM1F2-Ce3-FRB−tagged centriolar satellites (mean ± S.E.M; dark line, average; gray dashed line, a single individual cell; n = 14, 19, 17, 16, 14, and 9 from top to bottom; three to five independent experiments).

**Supplementary Figure 4.**
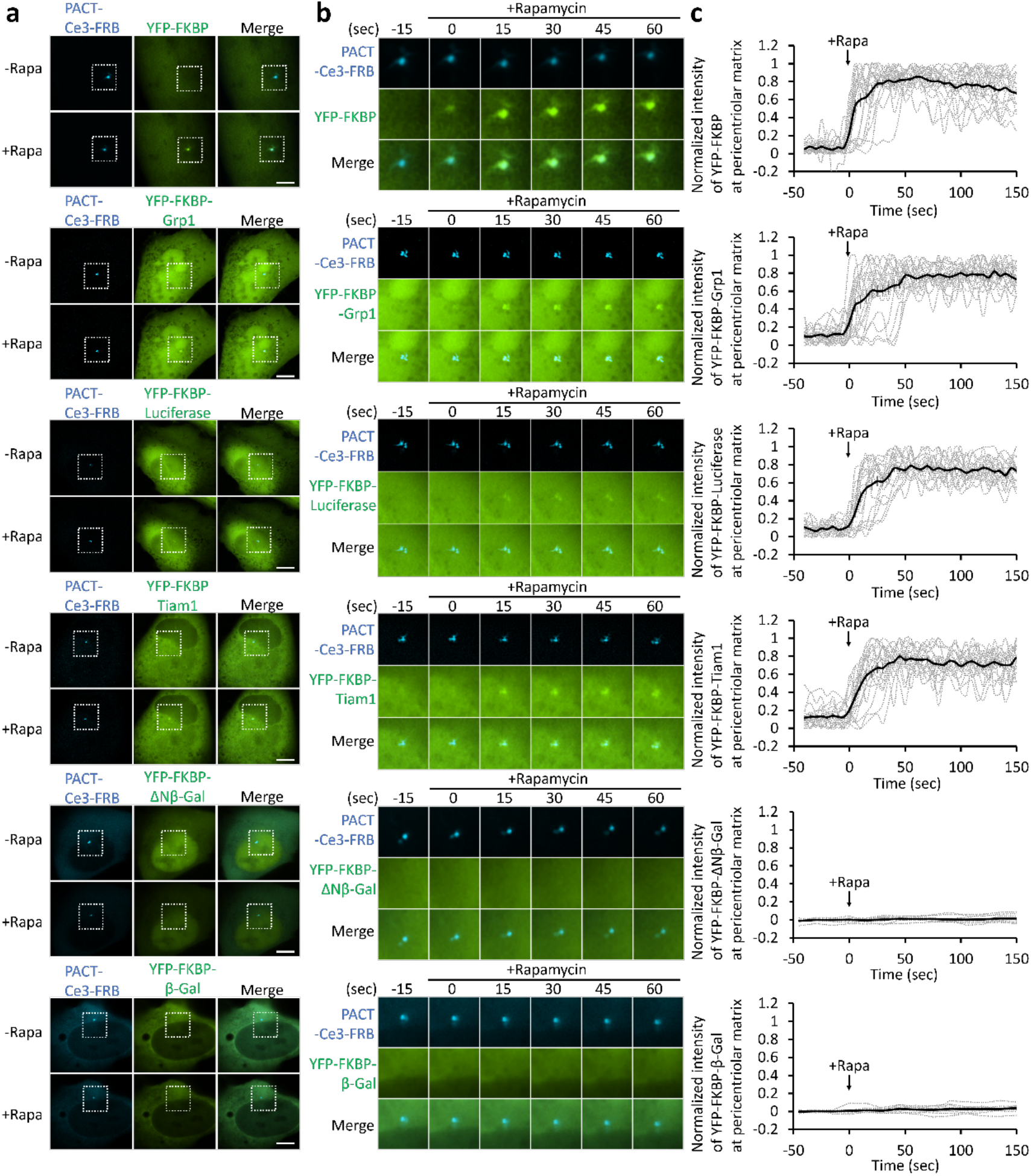
Accessibility of diffusion probes to the pericentriolar matrix. (a) U2Os cells co-transfected with PACT-Ce3-FRB and each YFP-FKBP−tagged varisized probe were treated with rapamycin (100 nM). The dotted line box indicates the pericentriolar matrix region. Scale bars: 10 µm. (b) Video frames from a movie of the pericentriolar matrix regions indicated by the dotted line boxes in A upon rapamycin treatment. (c) The normalized intensity of YFP-FKBP−tagged diffusive probes at the PACT-Ce3-FRB−tagged pericentriolar matrix (mean ± S.E.M.; dark line, average; gray dashed line, a single individual cell; n = 20, 19, 18, 18, 10, and 13 cells from top to bottom; three to five independent experiments).

**Supplementary Figure 5.**
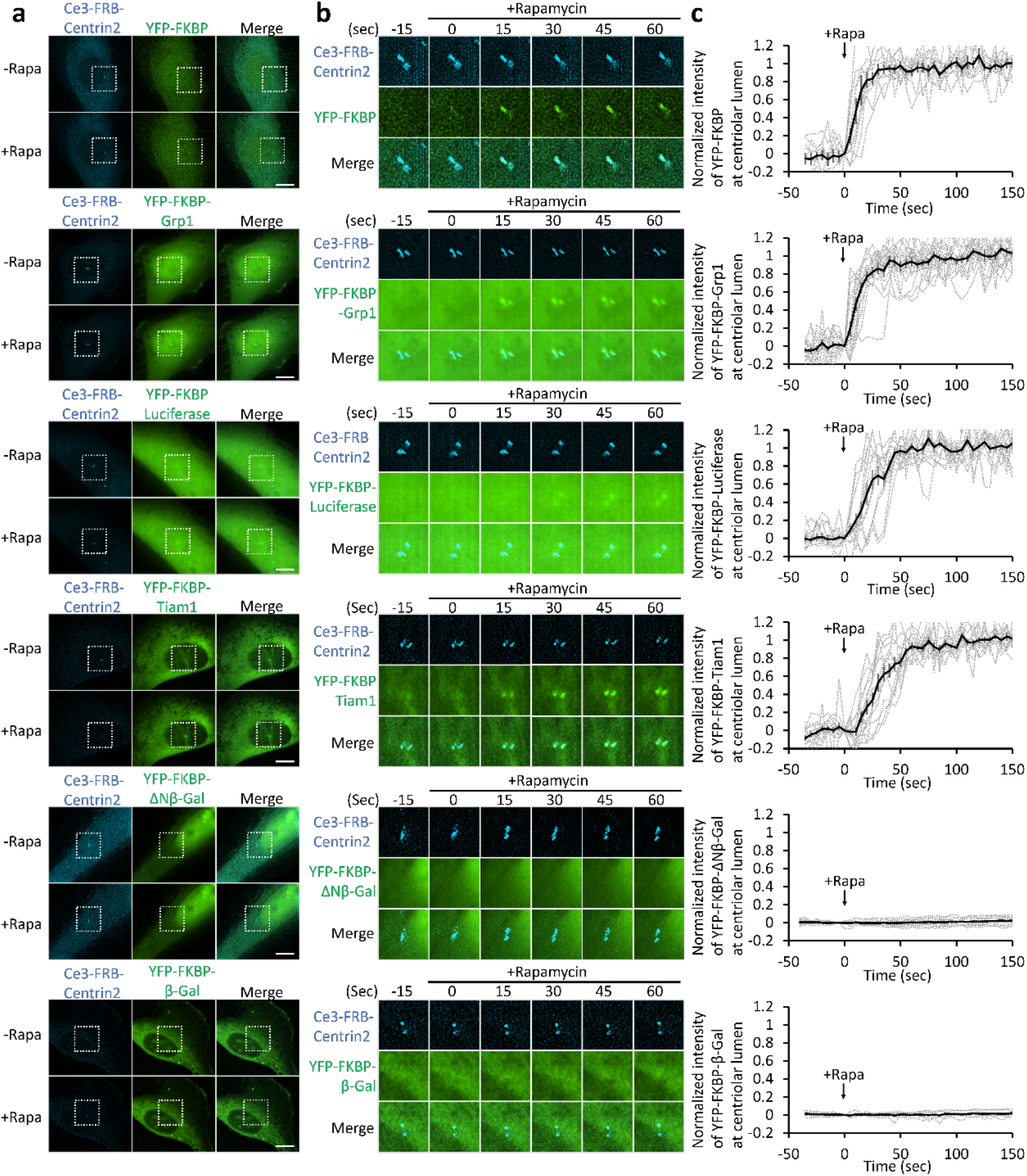
Accessibility of diffusion probes at centrioles. (a) U2Os cells co-transfected with Ce3-FRB-Centrin2 and each YFP-FKBP−tagged varisized probe were treated with rapamycin (100 nM). The dotted line box indicates the centriole region. Scale bars: 10 µm. (b) Video frames from a movie of centriole regions indicated by the dotted line boxes in A upon rapamycin treatment. (c) The normalized intensity of YFP-FKBP−tagged diffusive probes at Ce3-FRB-Centrin2−tagged centrioles (mean ± SEM; dark line, average; gray dashed line, a single individual cell; n = 10, 16, 13, 14, 13, and 9 cells from top to bottom; three to five independent experiments).

**Supplementary Figure 6.**
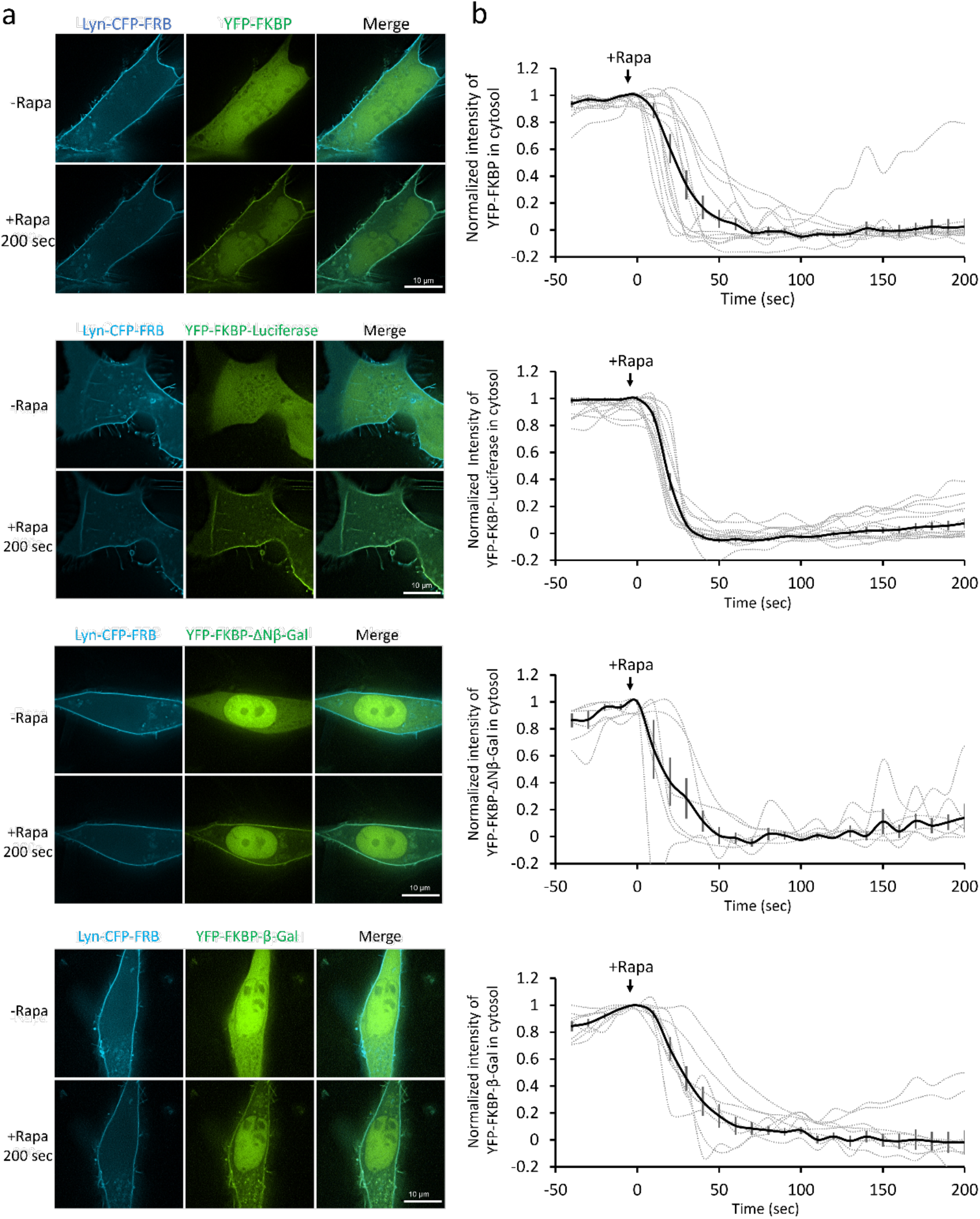
Translocation of YFP-FKBP−tagged diffusive probes onto the plasma membrane. (a) U2Os cells co-transfected with Lyn-CFP-FRB and YFP-FKBP, YFP-FKBP-Luciferase, YFP-FKBP-ΔNβ-Gal, or YFP-FKBP-β-Gal diffusive probes were treated with rapamycin (100 nM). Scale bar: 10 µm. (b) The normalized intensity of YFP-FKBP−tagged diffusive probes in the cytosol upon rapamycin treatment (mean ± SEM; dark line, average; gray dashed line, individual cell; n = 12, 14, 6, and 8 cells from top to bottom from two, three, three, and two independent experiments, respectively).

**Supplementary Video 1. Accessibility of diffusion probes to centriolar satellites**. U2Os cells co-transfected with centriolar satellites marker (PCM1F2-Ce3-FRB) and the indicated YFP-tagged-diffusion probes ranging from Rs of 3.2-7.6 nm were treated with rapamycin (100 nM) and imaged. Images were taken every 5 sec for 240 sec. Scale bar, 2.5 µm. See also Fig. 3 and supplementary Fig. 3.

**Supplementary Video 2. Accessibility of diffusion probes to pericentriolar matrix**. U2Os cells co-transfected with pericentriolar matrix marker (PACT-Ce3-FRB) and the indicated YFP-tagged-diffusion probes ranging from Rs of 3.2-7.6 nm were treated with rapamycin (100 nM) and imaged. Images were taken every 5 sec for 195 sec. Scale bar, 1 µm. See also Fig. 3 and supplementary Fig. 4.

**Supplementary Video 3. Accessibility of diffusion probes to centrioles**. U2Os cells co-transfected with centriole marker (Ce3-FRB-Centrin2) and the indicated YFP-tagged-diffusion probes ranging from Rs of 3.2-7.6 nm were treated with rapamycin (100 nM) and imaged. Images were taken every 5 sec for 240 sec. Scale bar, 1 µm. See also Fig. 3 and supplementary Fig. 5.

**Supplementary Video 4. The permeability of centrosomal diffusion barriers decreases in anaphase**. Hela cells co-transfected with PACT-Ce3-FRB and YFP-FKBP-Luciferase in interphase (top) or synchronized in anaphase (bottom) were treated with rapamycin (100 nM). Images were taken every 5 sec for 295 sec. Scale bar, 5 µm. See also Fig. 5.

